# The Unexpected Membrane Targeting of Marchantia Short PIN Auxin Exporters Illuminates Sequence Determinants and Evolutionary Significance

**DOI:** 10.1101/2024.04.29.591616

**Authors:** Han Tang, Adrijana Smoljan, Minxia Zou, Yuzhou Zhang, Kuan-Ju Lu, Jiří Friml

## Abstract

The plant hormone auxin and its directional transport are crucial for growth and development. PIN auxin transporters, on account of their polarized distribution, are instrumental in guiding auxin flow across tissues. Based on protein length and subcellular localization, the *PIN* family is classified into two groups: plasma membrane (PM)-localized long PINs and endoplasmic reticulum (ER)-localized short PINs. The origin of *PIN*s was traced to the alga *Klebsormidium*, with a single PM-localized long KfPIN. Bryophytes, the earliest land plant clade, represent the initial clade harboring the short PINs. We tracked the evolutionary trajectory of the short *PIN*s and explored their function and localization in the model bryophyte *Marchantia polymorpha*, which carries four short and one long PIN. Our findings reveal that all short MpPINs can export auxin, and they are all PM-localized with MpPINX and MpPINW exhibiting asymmetric distribution. We identified a unique miniW domain within the MpPINW hydrophilic loop region, which is sufficient for its PM localization. Phosphorylation site mutations within the miniW domain abolish the PM localization. These findings not only identify the essential sequence determinant of PINs’ PM localization but also provide a unique insight into the evolution of ER-localized PINs. Short MpPINW, which is evolutionarily positioned between the ancestral long PINs and contemporary short PINs, still preserves the critical region essential for its PM localization. We propose that throughout land plant evolution, the unique miniW domain has been gradually lost thus converting the PM-localized short PINs in bryophytes to ER-localized short PINs in angiosperms.

**IMPORTANT:**

- Manuscripts submitted to Review Commons are peer reviewed in a journal-agnostic way.
- Upon transfer of the peer reviewed preprint to a journal, the referee reports will be available in full to the handling editor.
- The identity of the referees will NOT be communicated to the authors unless the reviewers choose to sign their report.
- The identity of the referee will be confidentially disclosed to any affiliate journals to which the manuscript is transferred.

**GUIDELINES:**

- For reviewers: https://www.reviewcommons.org/reviewers
- For authors: https://www.reviewcommons.org/authors

**CONTACT:** The Review Commons office can be contacted directly at

## Introduction

Auxin, an essential plant hormone and morphogen, plays a crucial role during various stages of plant development throughout the entire plant’s life cycle (Gomes & Scortecci, 2021; Vanneste & Friml, 2009). Its key function operated via polar auxin transport (PAT) is to establish a gradient that accommodates the different needs at appropriate developmental stages (Friml & Palme, 2002). PAT is responsible for initial organ patterning, tissue development, and tropic responses (Han *et al*, 2021; Semeradova *et al*, 2020). This transport process is facilitated by a range of auxin transporters located on the plasma membrane (PM) (Naramoto, 2017). Among these transporters, a family called PIN-FORMED (PIN) proteins play a prominent role in exporting auxin from cells to drive PAT. The polar distribution of PIN proteins thus guides the direction of PAT toward specific target tissues (Adamowski & Friml, 2015).

The PIN family has undergone systematic examination regarding their polarization and role in auxin export during evolution (Tan *et al*, 2021). Typically, PIN proteins consist of two transmembrane domains (TMD) at the N-terminal and C-terminal, separated by a hydrophilic loop (HL). In Arabidopsis, PINs can be classified into two groups based on their subcellular localization and polypeptide length: PM-localized long PINs e.g. AtPIN1, 2, 3, 4, 7, and endoplasmic reticulum (ER)-localized short PINs e.g. AtPIN5, 6, 8 (Křeček *et al*, 2009; Mravec *et al*, 2009). The long and short PINs are also known as canonical long PINs and noncanonical short PINs, respectively. In PM-localized long PINs, multiple phosphorylation sites have been identified in the HL, where the phosphorylation status determines long PINs’ subcellular localization and polarization (Bennett *et al*, 2014; Křeček *et al*., 2009). In contrast, short PINs composed of two TMDs with shorter HL regions are predominantly situated in the ER membrane. Intriguingly, typical short PINs, AtPIN5 and AtPIN8, exhibit opposite orientations on the ER membrane, suggesting antagonistic functions contribute to auxin homeostatic maintenance within a cell (Ding *et al*, 2012).

Compared to long PINs, short PINs have received significantly less attention with focus on Arabidopsis. In recent studies, PIN genes from streptophytic algae *Klebsormidium flaccidum*, early land plants *Physcomitrium patens (P. patens)*, and *Marchantia polymorpha (M. polymorpha)* have been identified and partially characterized (Sisi & Růžička, 2020; Skokan *et al*, 2019). The sole PIN with long PIN characteristics, which is capable of auxin export, was found in Klebsormidium and localized to the PM in both Klebsormidium and Arabidopsis (Skokan *et al*., 2019). The ancestral PIN is therefore considered to be the long PIN. Bryophytes thus represent the initial clade of land plants having short PIN genes. In *P. patens*, only one short PIN was identified to localize to the ER (Viaene *et al*, 2014). In *M. polymorpha*, on the other hand, harbors four putative short PINs presumed to be ER-localized (Sisi & Růžička, 2020). The identification of short PINs in bryophytes suggests that short PINs evolved within this clade, possibly originating from the ancient KfPIN in Klebsormidium. However, the progress of the evolutionary transition from long PINs to short PINs remains elusive.

To unravel the evolutionary trajectory of the short PIN clade, we explored the functional aspects and intracellular localization of short PINs in Marchantia. Our findings reveal that all Marchantia short PINs can export auxin and induce growth phenotypes upon overexpression. Surprisingly, all of Marchantia’s short PINs were localized to the PM, with MpPINX and MpPINW displaying asymmetric localization. To uncover the crucial determinant for their PM and ER localization, we conducted bioinformatic analysis and identified a miniW domain with two critical phosphorylation sites within the HL of MpPINW, which is indispensable for MpPINW PM localization. Our results support that in Marchantia, short PINs still retain critical regions accounting for PINs’ PM localization. Furthermore, throughout land plant evolution, the critical region might have been gradually lost and evolved to ER-localized short PINs in angiosperms. Our research thus marks the transitional position of Marchantia short PINs between ancestral long PINs and contemporary short PINs along PINs’ evolutionary path.

## Results and Discussion

### Early-diverged Marchantia short PINs can export auxin

*Marchantia polymorpha (M. polymorpha)* genome encodes four short auxin efflux carriers MpPINW, MpPINX, MpPINV, and MpPINY. To position these genes within the context of other plants we performed phylogenetic analysis of short PINs among four species: two early divergent land plants *M. polymorpha* and *Physcomitrium patens*, the basal angiosperm *Amborella trichopoda*, and the model angiosperm *Arabidopsis thaliana*. The phylogenetic tree was generated by end-joined modeling in MEGA-X software. The unrooted result revealed that MpPINW has diverged from others at an early point, and Arabidopsis PINs are distant from bryophytic PINs. Amborella short PINs distribute evenly in the tree, agreeing with its intermediate position between bryophytes and Arabidopsis. (Figure 1A). Besides MpPINW, other MpPINs are closer related to PpPIND, which is localized to the endoplasmic reticulum (ER)(Viaene *et al*., 2014). The function and subcellular localization of *Marchantia* short PINs are reported below.

**Figure 1.**
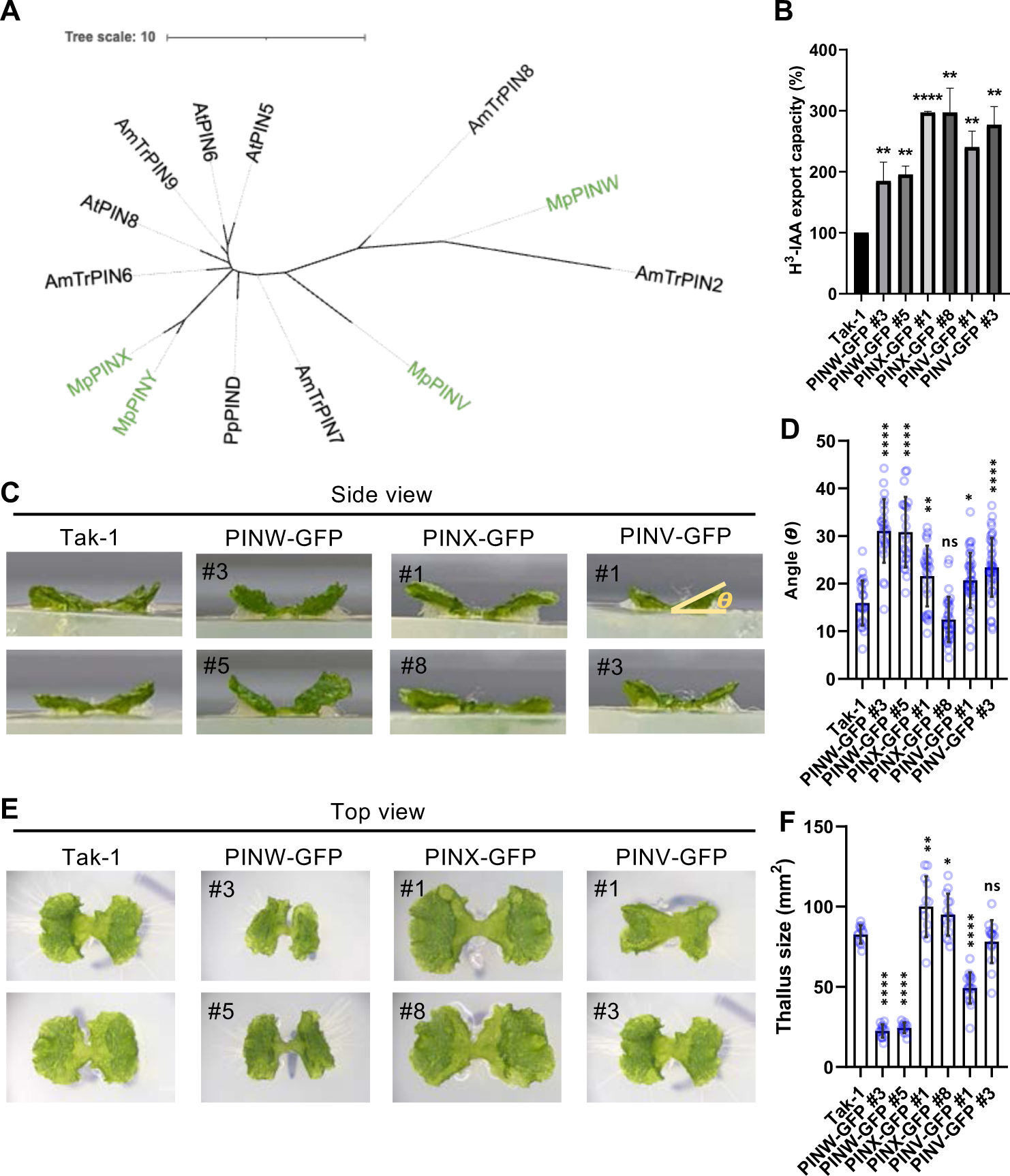
Early diverged Marchantia short PINs possess auxin export activity. A. Phylogenetic analysis of short *PINs* obtained from four selected species including two representative bryophytes *Physcomitrilum patens, Marchantia polymorpha*, two angiosperms *Amborella trichopoda*, and *Arabidopsis thaliana*. End-joined modeling was applied in the MEGA-X software. B. H^3^-IAA export assay was performed with the Marchantia overexpression lines as indicated. Transgenic plants were treated with H^3^-labeled IAA and recovered on H^3^-IAA-free medium for 24 hours. The exported H^3^-IAA in a liquid medium was detected as described previously (Lewis & Muday, 2009). Ten to fifteen 10-day-old Marchantia gemmae were used for one measurement, the graph shows the mean ± SD from three independent experiments. C. Vertical growth of the thallus when overexpressing MpPIN-GFP in Marchantia. Two independent lines labeled with numbers were selected for each transgenic line as indicated. Tak-1, the wildtype Marchantia served as control. D. Quantitative analysis of the vertical growth by manual angle measurement. The angle (8) between horizontal agar and the growth direction of the thallus was measured. Each blue circle represents a single measurement. 40>n>21 for each line. *** *P*<0.001, ** *P*<0.01, * *P*<0.05, Student’s *t*-test. E. The growth phenotype of the thallus in the same transgenic lines as seen in panel C. F. Quantitative analysis of the thallus growth. The thallus size was manually measured and corrected with *cosin8* based on the angle of vertical growth. The function of the magic wand and size measurement in the Fiji software were applied. Each blue circle represents a single measurement. 20>n>10 for each line. *** *P*<0.001, ** *P*<0.01, * *P*<0.05, Student’s *t*-test.

The Auxin transport ability of *M. polymorpha* short PINs was tested in overexpressing stable MpPIN-GFP lines that we generated for MpPINW-GFP, MpPINX-GFP, and MpPINV-GFP driven by CaMV *35S* promoter. As MpPINY is not expressed in all tissues it was not included in this study(Kawamura *et al*, 2022). To perform the auxin export assay, transgenic plants were cultivated in a liquid medium containing radioactive auxin (H^3^-IAA), followed by washing and one-day cultivation in a fresh medium. The level of H^3^-IAA exported to the fresh medium was then measured as previously described (Lewis & Muday, 2009). The wild-type Marchantia served as an internal control. All Marchantia short PINs were capable of exporting auxin. Although MpPINW showed lower export capacity compared to MpPINX and MpPINV (Fig 1B), overexpression of MpPINW-GFP resembled the phenotype observed when overexpressing Marchantia long PIN, MpPINZ-GFP, showing vertical growth and smaller size in thallus tissue (Tang *et al*, 2024) (Figure 1C-1F). In contrast, overexpression of MpPINX-GFP and MpPINV-GFP presented subtle phenotypes in the vertical growth and thallus growth (Fig 1C-1F). As the auxin export capacity was not fully reflected in the growth phenotypes, we reasoned that in addition to auxin export, overexpression of MpPINW-GFP may indirectly cause the phenotypes via yet unknown mechanisms. Our results demonstrate that all Marchantia short PINs are functionally conserved for auxin export.

### Marchantia short PINs are localized to the PM with asymmetric distribution

Based on the functional analysis, subcellular localization to the PM was anticipated. To prove this, the same MpPIN-GFP lines described above were examined. The MpPINW-GFP was localized to the PM with an unexpected asymmetric distribution along the PM in both rhizoid precursor cells and epidermal cells (Figure 2A, D-E). While PM localization was observed in every single line, the asymmetric distribution was shown in a small portion of cells of a gemma (Figure 2A). MpPINX-GFP showed similar localization, but the asymmetric distribution was not as profound as for MpPINW-GFP (Fig. 2B, 2E). The cells with asymmetrically distributed signals were selected for the quantification of the distribution index, which represents the degree of asymmetry in terms of signal distribution. To quantify the distribution index, we measured the signal intensity along the target PM in Fiji (Ferreira & Rasband, 2012). For a cell with asymmetrically distributed signals, the PM with the highest intensity was measured and divided by the signal measured on its opposite side. We reasoned that if the signals are equally distributed on the PM, the distribution index would be close to one, otherwise, the index would be larger than one if the signals are locally enriched (Fig. 2E, 2F). The quantification presented the asymmetric distribution of MpPINW-GFP and MpPINX-GFP, while MpPINV-GFP showed a ratio close to one, indicating an equal distribution on the PM (Fig. 2E). Whether this asymmetric distribution relates to their biological functions and may follow developmental cues, need to be further investigated.

**Figure 2.**
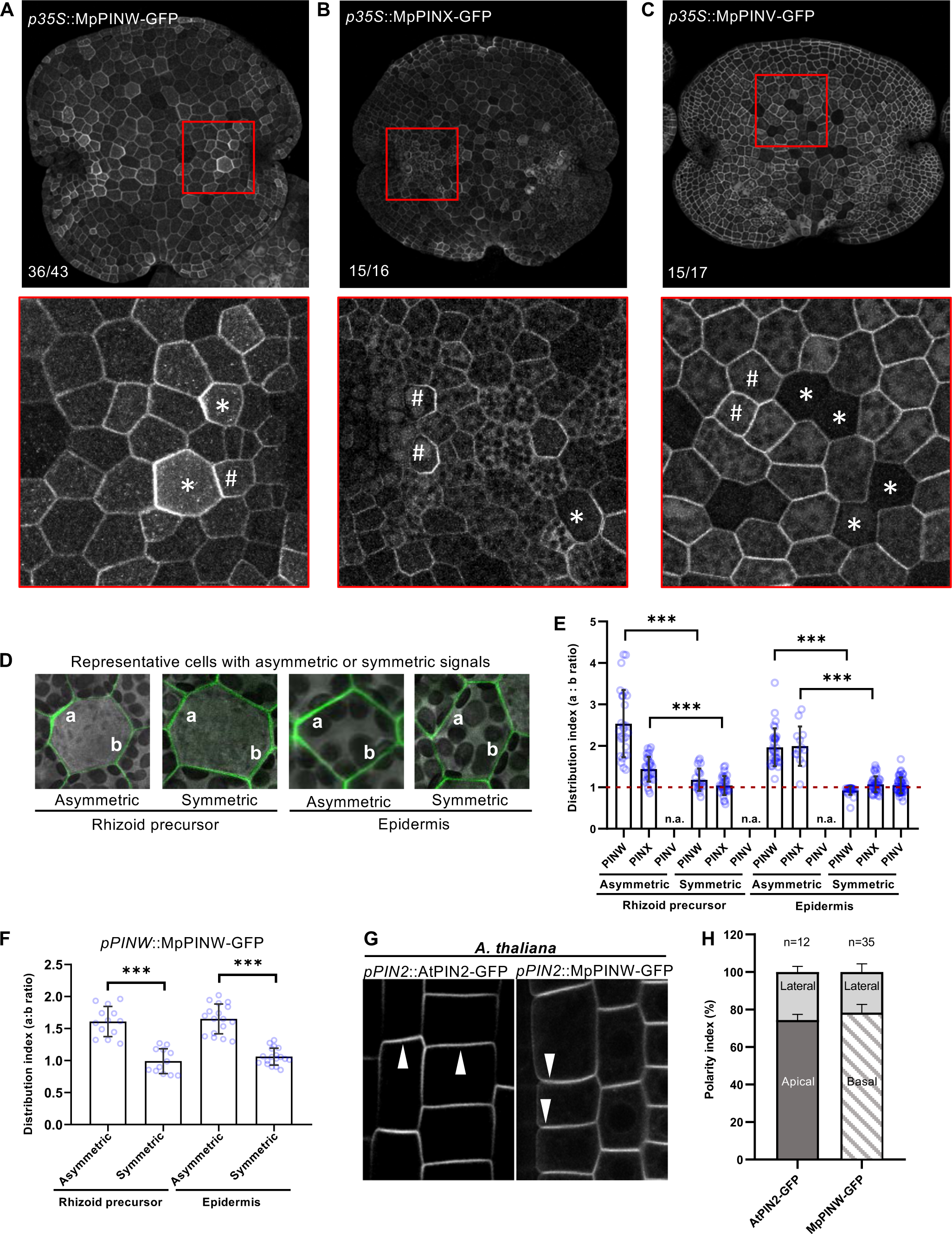
All Marchantia short PINs are localized to the plasma membrane with MpPINX and MpPINW exhibiting asymmetric distribution. A-C. Subcellular localization of MpPINW-GFP, MpPINX-GFP, and MpPINV-GFP in representative gemmae. The numbers labeled in lower left corners indicate the gemmae number that showed consistent signals as representative images over the total number of observed gemmae. The plasma membrane localization and asymmetric distribution are magnified in lower panels of A to C. Rhizoid precursor cells are indicated by asterisks, and epidermal cells are indicated by hashes. D. Representative cells display symmetric and asymmetric distribution of signals in rhizoid precursor cells and epidermal cells of MpPINW-GFP. Letters a and b point to the selected position of PM for the intensity measurement, applied to the quantification of the distribution index shown in E. E. Distribution index presents the occurrence of indicated MpPIN-GFP with symmetric or asymmetric signals in different tissues. The distribution index was calculated as ratio of signal intensity between “a” and “b” as shown in D. Each blue circle represents a single measurement. 30>n>12 for each line. *** *P<0.001,* Student’s *t*-test. F. Distribution index, quantified as described for E. Endogenously expressed MpPINW-GFP is asymmetrically localized to the plasma membrane in rhizoid precursor cells and epidermal cells. n>13 from three independent lines. *\*\*\* P<0.001,* Student’s *t*-test. G. AtPIN2-GFP and MpPINW-GFP driven by *AtPIN2* promoter exhibit apical and basal polarization in epidermal cells of Arabidopsis root. White arrowheads indicate the polar localization. H. Polarity index of AtPIN2-GFP and MpPINW-GFP in epidermal cells was quantified by dividing the polar signal by the signal on the lateral side of the same cell, as described previously(Zhang *et al*, 2019b).

Even under the same promoter control, MpPINV-GFP was exclusively expressed in epidermal cells, where it was evenly distributed on the PM (Fig. 2C and 2E). This result makes it unlikely that overexpression is causing the asymmetric distribution of MpPINW-GFP and MpPINX-GFP, but rather to the action of internal proteins. To further support this, we generated transgenic plants carrying *MpPINW-GFP* driven by its endogenous promoter. These constructs also showed an asymmetric distribution along the PM of epidermal cells and rhizoid precursor cells (Fig. S1, 2F). To test whether asymmetric localization of MpPINW also occurs in other species, we investigated MpPINW localization in Arabidopsis by using the endogenous *AtPIN2* promoter to control MpPINW-GFP. Like in Marchantia, asymmetric distribution of MpPINW-GFP was observed in Arabidopsis epidermal cells. While AtPIN2-GFP exhibited apical polarization as reported previously (Wiśniewska *et al*, 2006), MpPINW-GFP was localized along the basal membrane (Fig. 2G-H). This result demonstrates that the asymmetric localization of MpPINW is consistent in different species, while the regulatory machinery is likely species-specific thereby leading to different polar localization between AtPIN2 and MpPINW.

Although, based on the length of the PIN transporter, Marchantia short PINs were assumed to be ER-localized (Bennett *et al*., 2014), our results demonstrate that MpPINW-GFP rather resides at the PM with asymmetric distribution in both Marchantia and Arabidopsis. This unexpected result may be attributed to MpPINW’s longer HL region compared to typical short PINs e.g. AtPIN5 and AtPIN8. Given the HL region plays an essential role in long PINs’ PM localization and polarization (Tan *et al*., 2021), we suspected that the HL region of MpPINW may retain unidentified signal sequences accounting for its PM trafficking and asymmetric distribution.

To be noted, the asymmetric distribution of MpPINX and MpPINW did not display specific directional patterns (i.e. from the central section of the gemma towards the outside). The relative positions of cells with asymmetrically distributed signals across the entire gemma did not demonstrate any discernible patterns and may be randomly distributed in a gemma. In contrast, in Arabidopsis roots, the polarized PIN proteins accumulated at upper or lower sites of different tissues, where they govern auxin flow thereby contributing to tropic growth and root development(Adamowski & Friml, 2015; Friml & Palme, 2002; Křeček *et al*., 2009). Whether structural differences in tissue profiles between Arabidopsis and Marchantia relate to distinct biological roles of the enriched MpPINX and MpPINW remains uncertain. Further investigations are imperative to unravel the functional role of MpPINX and MpPINW at the enriched site.

### Localization of MpPINW to the PM requires the miniW domain

To search for a region within MpPINW that may account for its PM localization, we aligned the amino acid sequence of MpPINW with Arabidopsis ER-localized short PINs, AtPIN5 and AtPIN8 (Figure 3A). We found a short, unique domain (K168-V223, hereafter named miniW domain) within the hydrophilic loop (HL) of MpPINW, which made it a potential targeting candidate (Fig. 3A). To test this, we generated a truncated MpPINW-miniWΔ-GFP line that carries MpPINW-GFP without the miniW sequence. Protoplast transformation together with the ER tracker staining verified its colocalization with the ER structure (Fig. S2A). The full-length MpPINW-GFP was localized to the PM, while the MpPINW-miniWΔ-GFP was mainly localized to the ER with high colocalization occurrence with the ER tracker (Fig. S2A).

**Figure 3.**
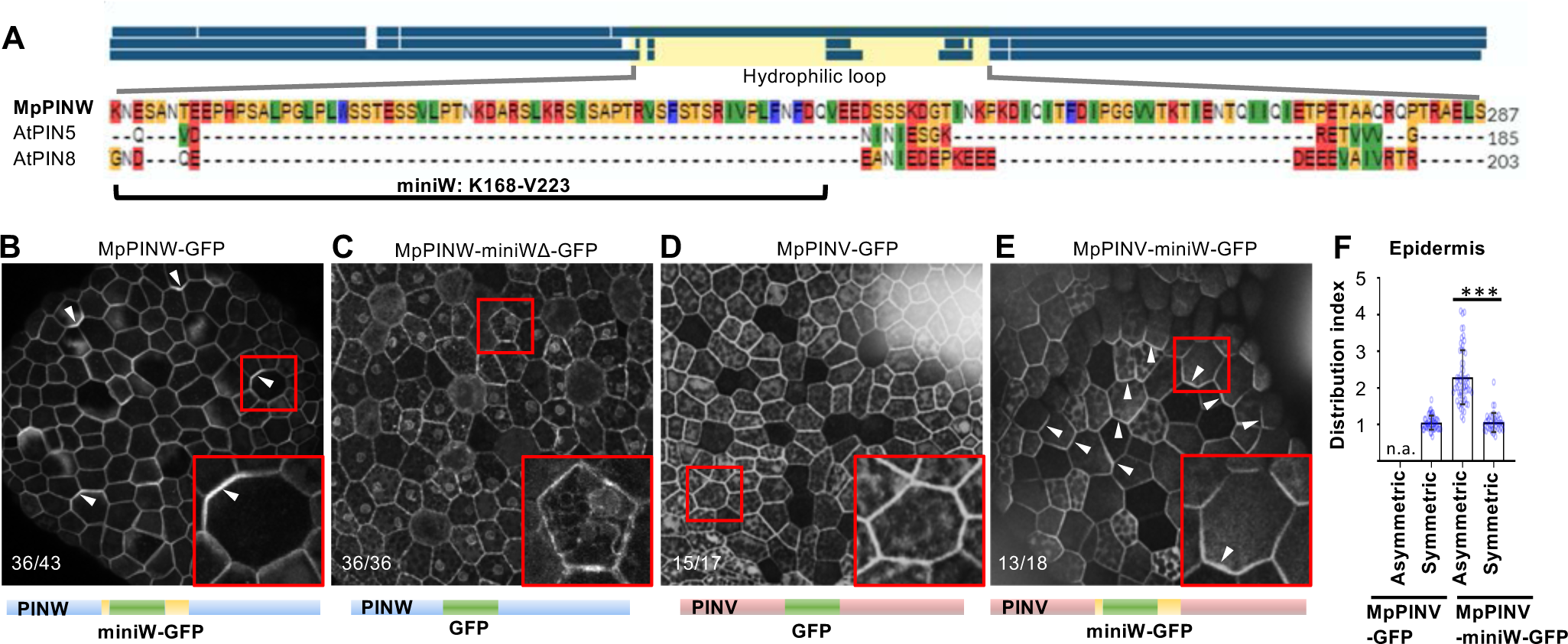
The miniW domain of MpPINW is essential for PINs’ asymmetric PM localization. A. Amino acid alignment of MpPINW, AtPIN5, and AtPIN8. An extra domain only exists in MpPINW and is named the miniW domain. B-C. Subcellular localization of full-length MpPINW-GFP and MpPINW-miniWΔ-GFP without the miniW domain. The MpPINW-miniWΔ-GFP was localized to the ER-like structure. The protein composition is presented below. D-E. Subcellular localization of full-length MpPINV-GFP and MpPINV-miniW-GFP with the miniW domain insertion. The protein composition is presented below each image. Plasma membrane-localized MpPINV-GFP shows asymmetric distribution with the miniW insertion. The occurrence was shown in the lower left corner. F. Distribution index shows the localization changes between symmetric-distributed MpPINV-GFP and asymmetric-distributed MpPINV-miniW-GFP in epidermal cells. Each blue circle represents one measurement. 59>n>40 for each line. *** *P<0.001,* Student’s *t*-test.

In Marchantia, while MpPINW-GFP predominantly localized to the PM, we found that MpPINW-miniWΔ-GFP mainly localized to the ER-like structure (Figure 3B-3C). In the root epidermal cells of Arabidopsis, the MpPINW-miniWΔ-GFP showed consistent localization change from PM to ER (Fig S2B). The only Marchantia short PIN that showed even distribution is MpPINV (Fig. 2C), which lacks the miniW region. Therefore, to test whether the miniW domain is sufficient to change the distribution of MpPINV-GFP, we inserted the miniW domain into the MpPINV coding sequence at the corresponding position. The signal of MpPINV-miniW-GFP was asymmetrically distributed on the PM (Fig. 3D, 3E), supported by the distribution index (Fig. 3F). These results validate the significance of the miniW domain for the asymmetric distribution of Marchantia short PINs.

How ER-localized short PINs evolved from the ancient PM-localized long KfPIN has been mysterious since decades (Bennett *et al*., 2014). The phylogenetic analysis revealed that *MpPINW* is the earliest diverged short *PIN* (Fig. 1A). We discovered that short MpPINW is localized to the PM and its miniW domain contributes to the PM-to-ER location transition and the asymmetric distribution (Fig. 2, 3). Thus, our results suggest that during land plant evolution, the *PIN* family may have gradually lost the miniW domain thereby shifting their PM localization to the ER, as contemporary short PINs present in Arabidopsis.

### Mutations at phosphorylation sites change MpPINW localization

In PM-localized long PINs, multiple phosphorylated residues have been identified, and the phosphorylation status of PINs is pivotal to their localization patterns in different tissues (Friml *et al*, 2004; Michniewicz *et al*, 2007; Tan *et al*., 2021; Zhang *et al*, 2010). It is feasible that regulation of MpPINW intracellular trafficking may share similar mechanisms to long PINs. We aligned the MpPINW amino acid sequence with canonical long PINs including MpPINZ, PpPINA, AtPIN1, and AtPIN2 (Fig. 4A). This allowed identification of putative phosphorylation sites within the miniW domain, S180, and S193, which have been predicted to be phosphorylated in Arabidopsis long PINs (Dory *et al*, 2018).

**Figure 4.**
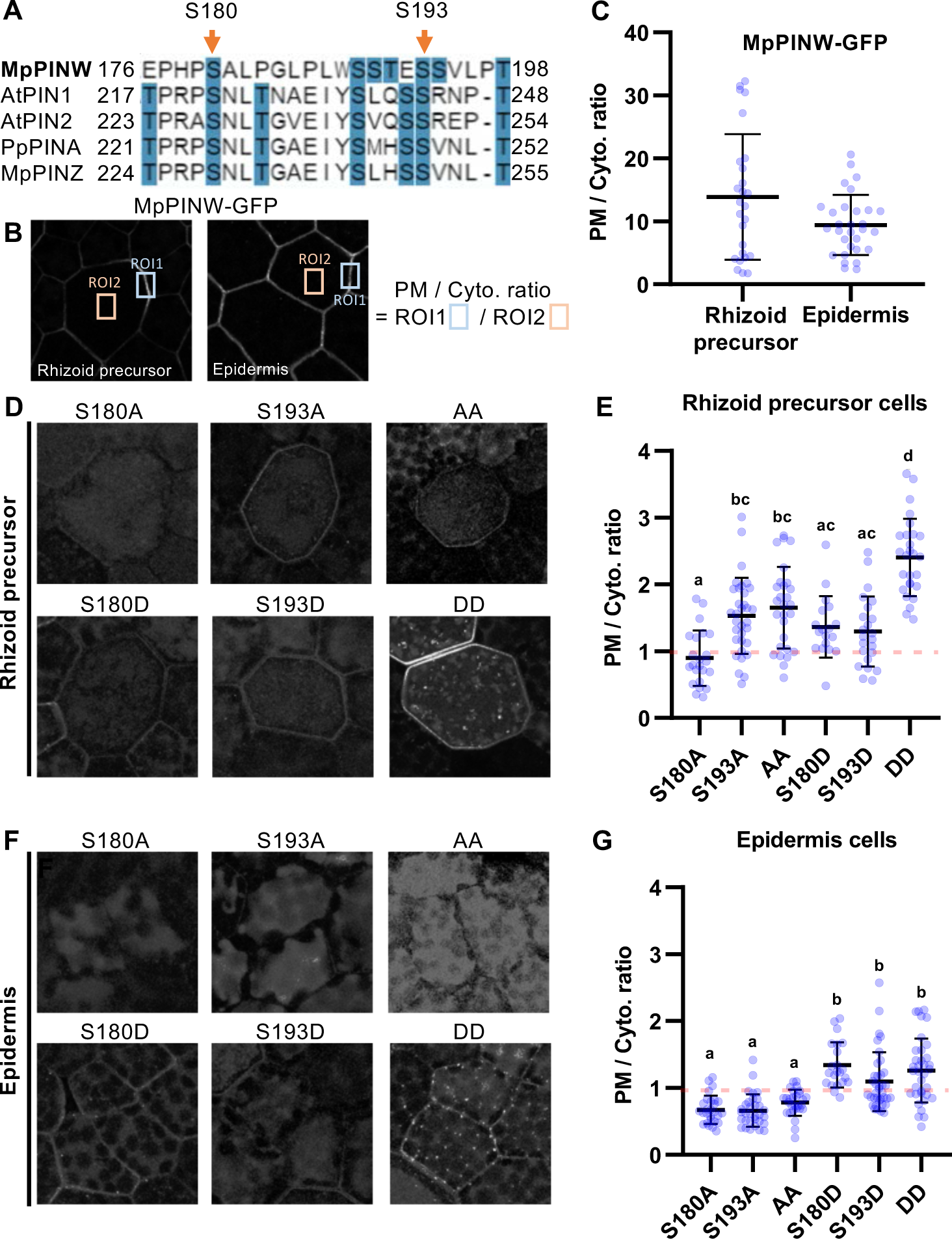
Mutations in putative phosphorylation sites within the miniW domain abolish MpPINW PM localization. A. Alignment of MpPINW and four long PINs as indicated with a focus on the miniW domain. Two putative phosphorylation sites are marked by orange arrows. B. Representative cells with MpPINW-GFP localized on the plasma membrane, and ROI 1 and ROI 2 represent the selected area for the ratio calculation. The intensity ratio is quantified by dividing the intensity of the intensity enriched area (ROI 1) on the plasma membrane (PM) by the intensity of the randomly selected ROI 2 in the cytoplasm (Cyto). C. The intensity ratio of MpPINW-GFP shows the majority of MpPINW-GFP are localized to the PM in both rhizoid precursor cells and epidermal cells. D. The subcellular localization in rhizoid precursor cells and E. The quantification analysis of the intensity presented in PM to cytoplasm ratio of mutated MpPINW-GFP as indicated. F. The subcellular localization in epidermal cells and G. the quantification analysis of the intensity presented in PM to cytoplasm ratio of mutated MpPINW-GFP as indicated. The letter above represents statistic groups with P<0.05 using ANOVA and Tukey’s HSD test.

We generated phosphor-mimic and phosphor-dead mutants to test if their phosphorylation status is critical for MpPINW PM localization. In rhizoid precursor cells, the S180A mutant showed almost no PM localized signal, rather most signals accumulated in the cytoplasm. In contrast, other single or double mutants presented localization in both PM and cytoplasm with a much lower PM-to-cytoplasm ratio compared to the intact MpPINW-GFP (Figure 4B-4E). Double site phosphor-mimic mutant showed a strong signal at PM, while the cytoplasmic signal was still higher than MpPINW-GFP. In epidermal cells, all phosphor-dead mutants showed exclusively cytosolic signals, whereas phosphor-mimic mutants displayed a weak PM signal in addition to cytosolic signal (Fig. 4F, 4G). Our results revealed that single mutations on the phosphorylation sites of MpPINW lead to localization changes in distinct tissues. It suggests that tissue-specific regulatory pathways may already exist in early divergent land plants.

## Conclusions

Short PINs of early diverging land plant Marchantia present unexpected PM localization with asymmetric localization patterns, for which the miniW domain of MpPINW plays an essential role, presumably via phosphorylation modifications. We propose that across land plant evolution, long PINs may progressively shorten their central, hydrophilic loop and finally lose the miniW domain, leading to a shift in localization from the PM to the ER. Here we identified an evolutionary intermediate—Marchantia short PINs—which may connect the ancestral PM-localized long PIN to contemporary ER-localized short PINs.

## Materials and Methods

### Plant materials and transformations

Throughout this research, we utilized *Marchantia polymorpha* Takaragaike-1 (Tak–1). The plants were cultured on 1/2 Gamborg B5 medium containing 0.8% Daishin agar in a 21°C growth chamber under 16/8 day/night cycles, with 60 µmol m^-2^s^-1^ photons from white LED lights.

Transgenic Marchantia plants were generated using the agrobacterium transformation method described previously(Kubota *et al*, 2013). Briefly, the apical meristematic region of each two-week-old thallus was excised, and the thallus was then divided into four pieces. After culturing on 1/2 B5 agar plates with 1% sucrose for three days, the cut thalli were transferred to 50 ml of 0M51C medium in 200 ml flasks supplemented with 200 µM acetosyringone (4’-Hydroxy-3’,5’-dimethoxyacetophenone). Agrobacteria harboring the target construct, with an OD600 of 1 density, were added for co-cultivation for an additional three days. Subsequently, the transformed thalli were washed and plated on 1/2 B5 plates with corresponding antibiotics for selection. Independent transformed lines (T1) were isolated, and the second generation (G1) from independent T1 lines was generated by sub-cultivating single gemmaling, which emerged asexually from a single initial cell(Shimamura, 2016). The subsequent generation (G2 generation) was used for analyses.

For Arabidopsis, seeds were surface sterilized and plated on to 1/2 MS medium with 1% agar plates. After 3 days of stratification, plates were vertically cultured in a 23°C growth chamber with 150 µmol m^-2^s^-1^ photons light intensity and 16/8 day/night cycles.

Arabidopsis transgenic lines were generated by the floral dipping method(Clough & Bent, 1998) with slight modification. The floral buds were dipped with a small amount of Agrobacterium by pipetting and kept in the dark for another 24 hours with high humidity.

### Microscopy

Confocal microscopy utilized the Leica Stellaris 8 system with hybrid single-molecule detectors (HyD) and an ultrashort pulsed white-light laser (WLL; 70%; 1 ps at 40 MHz frequency). Leica Application Suite X served as the software platform, with imaging conducted using an HC PL APO CS2 40x/1.20 water immersion objective. For GFP-containing images, a 488 nm white light laser was selected, with the detection range set between 500 nm to 525 nm. The following imaging settings were used: scan speed of 400 Hz, resolution of 1024 x 1024 pixels. A tau-gating model, capturing photons with a lifetime of 1.0-10.0 ns, was employed for all Marchantia imaging to mitigate autofluorescence.

For Arabidopsis root imaging, 4-day-old seedlings of each indicated genotype were used. Seedlings were mounted on a slice with growth medium and then placed into a chambered coverslip (Lab-Tek) for imaging. Imaging was conducted using a laser scanning confocal microscopy (Zeiss LSM800, 20x air lens), with the default setting for GFP detection applied.

For protoplast co-localization analysis, live-cell imaging was performed using a Leica SP8X-SMD confocal microscope equipped with HyD detectors and an ultrashort pulsed white-light laser (WLL 50%; 1 ps at 40 MHz frequency). Leica Application Suite X was used for microscope control, and an HC PL APO CS2 20x/1.20 water immersion objective was used for observing the samples. The following imaging settings were used: scan speed of 400 Hz, resolution of 1024 x 1024 pixels, and standard acquisition mode for the hybrid detector.

Surface area measurement of 14-day-old Marchantia gemmae was performed using a stereomicroscope (SZN71, LeWeng, Taiwan) with a CCD camera (Polychrome M, Liweng, Taiwan), and images were processed using Fiji (ImageJ, https://imagej.net/software/fiji/) software.

### Phenotyping

Gemmae were cultured on a 1/2 Gamborg B5 agar plate for 14 days. A rectangular agar cube with an individual plant on top was cut from the plate by a scalpel and transferred onto the center of a slide. The slide was positioned on the surface of a laminar flow at a fixed distance to the age, and the images were captured using a Google Pixel 8 cell phone camera. The angle was further measured using Fiji software.

### Plasmid construction

All constructs were performed using the Gateway^TM^ system (Invitrogen) as recommended by the user manual. The pMpGWB vectors, developed for Marchantia transformation (Ishizaki *et al*, 2015), were used as the final destination vectors.

*pDONR221-MpPINW-, MpPINX-, and MpPINV-GFP* were obtained from previously published plasmids (Zhang *et al*, 2019a). The fragments in the entry vector were transferred into the destination vector pMpGWB102, containing a *35S* promoter, using the LR Clonase^TM^ II enzyme according to the manual (Invitrogen) to generate p*35S::MpPINW*-, *MpPINX*- and *MpPINV-GFP*.

For endogenous *MpPINW* promoter-driven constructs, a 3.5k *MpPINW* promoter was amplified with primers listed in Table S1 and cloned into the pENTR^TM^/D-TOPO^TM^ vector as the manual suggested. The *MpPINW-GFP* fragment was amplified from the *pDONR221-MpPINW-GFP* using primers listed in Table S1, and the *MpPINW* promoter-containing entry vector was linearized by PCR with a back-to-back primer set targeting the end of the promoter region (listed in Table S1). The insert and linearized vector were fused using the seamless cloning (SLiCE) method (Zhang *et al*, 2014). The resulting *MpPINWpro::MpPINW-GFP* fragment was further transferred into pMpGWB101 vector by the LR Clonase^TM^ II enzyme according to the manual.

*MpPINW-ΔminiW-GFP* was generated by site-directed mutagenesis PCR with primers listed in Table S1 to exclude the miniW domain and the inserted GFP in the *pDONR221-MpPINW-GFP* vector. A second *GFP* fragment was amplified with the primers listed in Table S1 to fused with the linealized pDONR221-*MpPINWΔminiW* by SLiCE method. The *MpPINWΔminiW-GFP* fragment was then transferred into the pMpGWB102 vector with the LR reaction. The *PINW* single and double amino acid mutants were generated by site-directed mutagenesis PCR (primers listed in Table S1) with the *pDONR221-MpPINW-GFP* vector, and the mutated fragments were transferred into the pMpGWB102 vector using the LR reaction.

For *PIN2pro::PIN2-* and *MpPINW-GFP* Arabidopsis lines, plants were generated in a previous publication in the group(Zhang *et al*., 2019a).

### Phylogenetic analysis

The phylogenetic analysis for full-length amino acid sequences of all examined PINs was carried out in MEGA X program(Kumar *et al*, 2018) and the evolutionary history was inferred by using the end-joined modeling and JTT matrix-based model with default settings(Jones *et al*, 1992). The results were imported into iTOL (https://itol.embl.de/) for visual illustration. The alignment and identity index were produced by an online CLUSTAL alignment program with default settings.

### *A*uxin export assay

The auxin export assay performed with transgenic Marchantia plants was modified based on the protocol developed for Arabidopsis seedlings(Lewis & Muday, 2009). In brief, around fifteen 10-days-old gemmae were transferred to a liquid growth medium for another 3 days with gentle shaking, followed by 10 nM ^3^H-IAA treatment for 24 hours. The radioactive tissues were then washed twice with sterile H_2_O, and were cultivated in fresh growth medium for another 24 hours. The cultivated medium was then collected to mix with ScintiVerse BD cocktail (Fisher, SX18-4) in 1:30 (v:v), and the export of auxin was measured by the Scintillation counter (Beckman, LS6500).

### Distribution index analysis

The PIN-GFP signals on the PM were quantified and presented as a distribution index. For asymmetric distribution, the intensity of the strongest signal on the PM (intensity a) was measured by the line tool in Fiji with a 3-pixel thickness. The intensity of the opposite side (intensity b) was measured in the same way. The intensity b was then divided by the intensity a as the distribution index. The index is close to 1 indicating the symmetry of signals, while larger than 1 indicates the asymmetry of the protein distribution.

### Statistical analysis

For phenotype analysis (Figure 1D and 1F), Student’s t test was performed to compare *PINs* overexpression lines with wildtype. For distribution index analysis (Figure 2E, 2F and 3F), the means of groups with asymmetric and symmetric signals were compared and analyzed by Student’s t test as paired samples. For the intensity ratio in cytoplasm versus PM (Figure 4E and 4G), Student’s t test was performed to compare each pair of transgenic lines.

### Protoplast transformation

Protoplasts were prepared and transformed as previously described(Mathur & Koncz, 1998). Plasmids were prepared with an E.Z.N.A. Plasmid Maxi Kit I (Omega Bio-Tek). 10 micrograms of each plasmid was transformed into the protoplasts. The transformed protoplast cells were incubated in the dark at room temperature for 12 h to 16 h before imaging under an LSM800 confocal microscope (Zeiss).

## Acknowledgment

The authors sincerely thank Dr. Shutang Tan for experimental support, and Dr. Barbara Kloeckener Gruissem for critical reading and constructive advice on the manuscript. This work was supported by the European Research Council Advanced Grant (ETAP-742985 to H.T. and J.F.) and by the Ministry of Science and Technology (grant 112-2636-B-005-001-to K.-J.L.).

## Author contributions

H.T., K.-J.L., and J.F. initiated and designed the experiments. Y. Z. provided Arabidopsis transgenic line. A. S. and M. Z. performed the auxin export assay. K.-J.L. performed Marchantia experiments and H. T. carried out quantitative analysis and wrote the manuscript.

---

**Figure S1.**
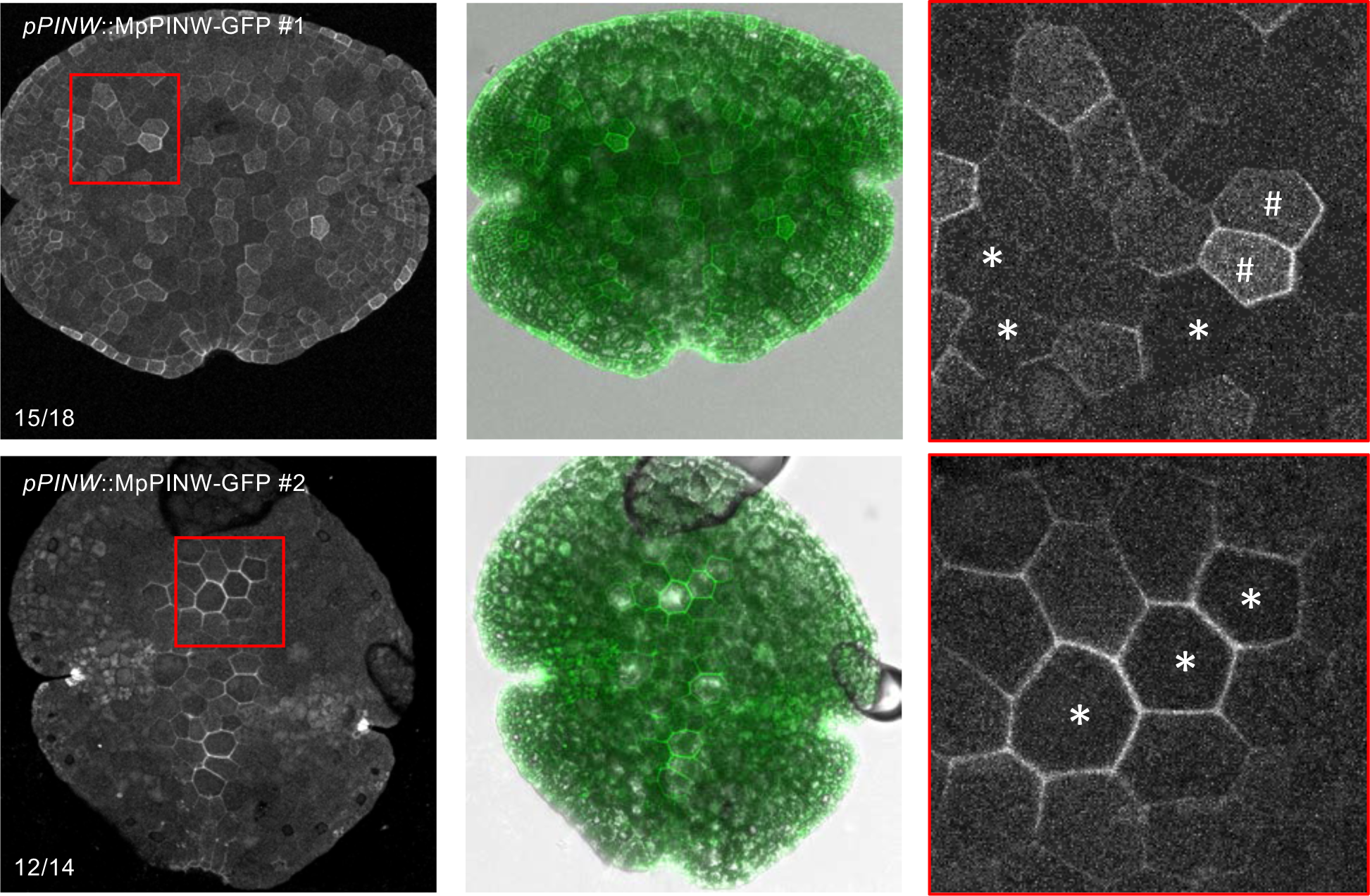
Endogenous PINW are asymmetrically distributed in different tissues. Subcellular localization of MpPINW-GFP, driven by its endogenous promoter, in representative gemmae. The number of gemmae that showed signals as the representative image compared to the total number of served gemmae is shown in the lower left corner. The plasma membrane localization and asymmetrictribution are magnified in the right panels. Rhizoid precursor cells are indicated by asterisks, and idermal cells are indicated by hashes.

**Figure S2.**
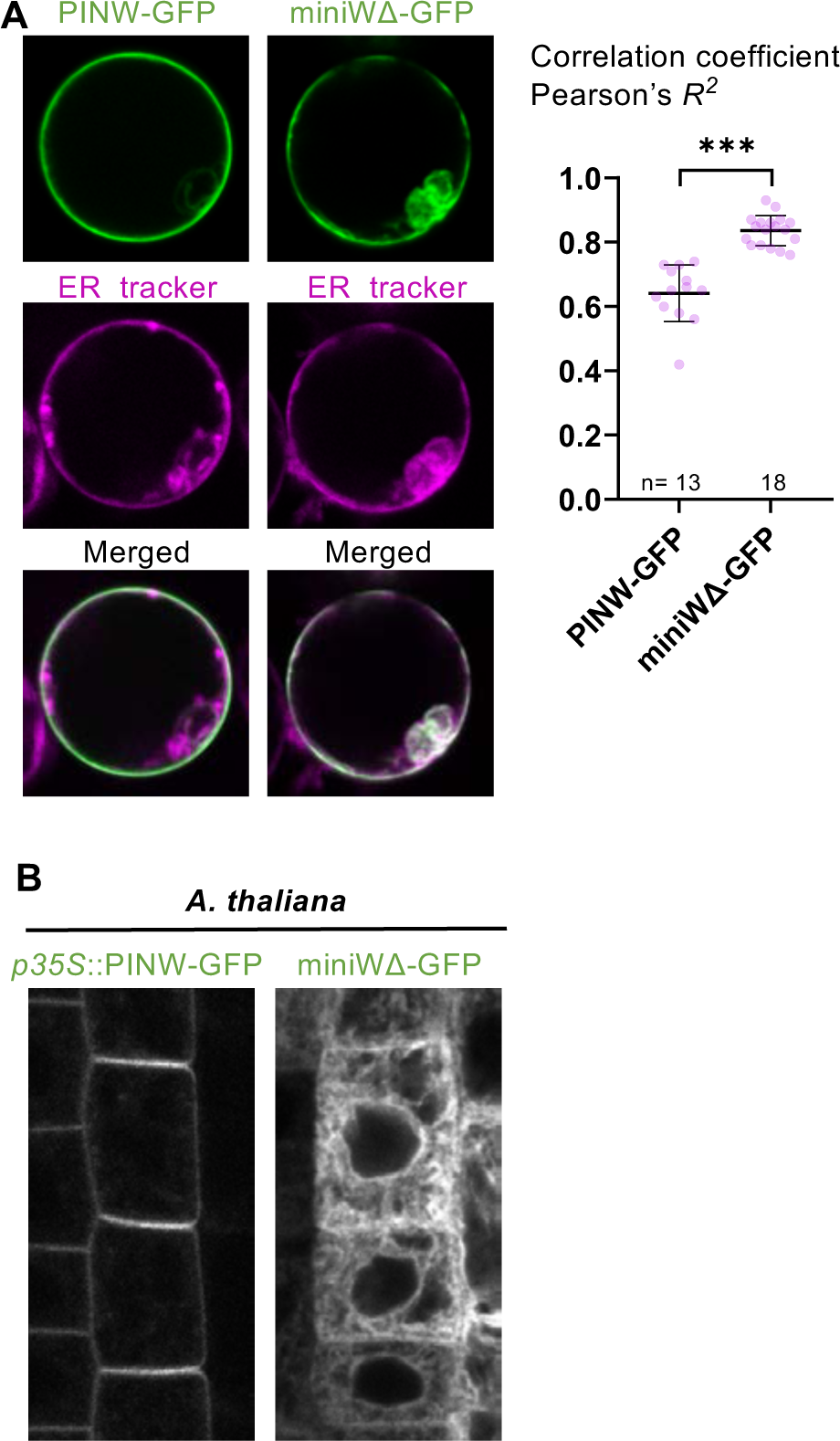
MpPINW colocalizes with ER in protoplast staining. A. Protoplast derived from Arabidopsis roots was transformed with *p35S*::MpPINW-GFP or *p35S*::MpPINW-miniW Δ -GFP, coupled with ER tracker staining colored in magenta. The Pearson’s correlation coefficient was analyzed by Fiji application. B. Arabidopsis transformants carrying the same constructs were observed. Intact MpPINW-GFP shows PM localization, while PINW-miniWΔ-GFP is translocated to ER-like structure.

